# Molecular structure and nickel binding capacity of *Proteus mirabilis* UreE

**DOI:** 10.64898/2025.12.19.695641

**Authors:** Jiayi Pan, Sarah L. Mueller, Nuren Tasneem, Yang Wu, Emily J. Furlong

## Abstract

UreE is a nickel chaperone required for the safe and efficient delivery of nickel to the active site of the metalloenzyme, urease; a key virulence factor of the urinary tract pathogen, *Proteus mirabilis*. We investigated the structural features of *P. mirabilis* UreE using protein X-ray crystallography and its nickel-binding capacity by inductively coupled plasma-mass spectrometry. Here, we report a 2.0 Å crystal structure of homodimeric *Pm*UreE and show it has capacity to bind five nickel ions per dimer. Truncation of the histidine-rich C-terminus reduced nickel binding capacity by two nickel ions per dimer and comparison with homologous UreE structures allowed the assignment of putative nickel binding sites within the *Pm*UreE structure. These findings increase our understanding of how *Pm*UreE binds nickel and ultimately prevents this toxic metal from causing significant cellular damage in *P. mirabilis*.

## 1. Introduction

*Proteus mirabilis* is a gram-negative bacterium and a leading contributor to catheter-associated urinary tract infections (CAUTIs), known for its ability to induce crystalline biofilms and formation of urinary stones (Armbruster *et al*., 2018, Armbruster *et al*., 2017). Crystalline biofilms and urinary stones block catheters but also protect *P. mirabilis* from the immune system and antibiotics, enabling persistent infection of the urinary tract (Stickler & Feneley, 2010, Sabbuba *et al*., 2004). The critical protein responsible for the crystal-forming trait of *P. mirabilis* is urease (Jones *et al*., 1990, Mobley & Hausinger, 1989). Urease is a nickel-dependent enzyme that hydrolyses urea and produces ammonia, which subsequently increases the pH of the local environment. In the urinary tract, this causes the formation of struvite and apatite crystals from the minerals found in urine (Schaffer *et al*., 2016, Li *et al*., 2002). Active urease is required for *P. mirabilis* pathogenesis; mutants that do not express the enzyme or are deficient in nickel import, cannot induce urinary stone formation and have significantly reduced fitness in a murine urinary tract infection (UTI) model (Jones *et al*., 1990, Brauer *et al*., 2020).

Despite its necessity for urease activity, nickel can also be toxic to bacteria at high intracellular concentrations, disrupting cellular homeostasis and inducing oxidative stress. Therefore, bacterial nickel uptake and distribution must be strictly controlled to prevent toxicity and ensure appropriate metal allocation to nickel-dependent enzymes (Zeer-Wanklyn & Zamble, 2017). Most research on the maturation processes of bacterial ureases has been performed in *Klebsiella aerogenes* and *Helicobacter pylori*. In these organisms, the nickel-chaperone, UreE, has an essential role in capturing nickel in the bacterial cytoplasm and delivering it to UreG, which then goes on to interact with UreD and UreF to deliver the toxic metal into the urease active site (Nim *et al*., 2023, Nim & Wong, 2019).

Structural and biochemical analysis of UreE from *H. pylori*, *K. aerogenes*, and *Sporosarcina pasteurii* have shown that it functions as a dimer (Remaut *et al*., 2001, Shi *et al*., 2010, Song *et al*., 2001, Banaszak *et al*., 2012, Zambelli *et al*., 2013). The core fold is conserved between these species and includes a GNRH motif that sits at the dimer interface, with the histidine residues from each protomer contributing to the coordination of one nickel ion. There are variations between UreE homologues, primarily in the C-terminus, which often features a histidine-rich tail with variable length and amino acid composition (Fig. S1). As a result of this, the number of nickel ions bound by UreE homologues and the nickel binding sites also vary. *H. pylori* only has one histidine in its C-terminal region and studies report an overall nickel capacity of ∼1 Ni^2+^ per dimer, which is coordinated between the histidine residues at the dimer interface and one C-terminal histidine (Bellucci *et al*., 2009, Shi *et al*., 2010, Banaszak *et al*., 2012). *S. pasteurii*, has two C-terminal histidine residues and isothermal titration calorimetry and crystallography results showed 2 Ni^2+^ binding per dimer with positive cooperativity between the two sites (Stola *et al*., 2006). The high-affinity site involves the conserved dimer interface site and one C-terminal *Sp*His145, while the second site involves *Sp*His145 and *Sp*His147 from the other protomer (Stola *et al*., 2006, Zambelli *et al*., 2013). *K. aerogenes* UreE has 10 C-terminal histidine residues and binds approximately 6 Ni^2+^ per dimer, whereas a C-terminally truncated variant (*Ka*H144*UreE, lacking the His-rich tail) binds only 2 Ni^2+^ per dimer (Lee *et al*., 1993, Brayman & Hausinger, 1996). The crystal structure of *Ka*H144*UreE however, supports the binding of 3 metal ions, suggesting the earlier experiments may underestimate nickel binding capacity (Song *et al*., 2001). The *Ka*H144*UreE variant remains functional for urease activation, indicating that the core dimer can support the delivery of Ni^2+^ to urease via UreG without the C-terminal histidine residues, albeit with slightly less efficiency (Brayman & Hausinger, 1996). Further mutational studies in *Ka*UreE identified another Ni^2+^ binding site within the core fold of each UreE protomer composed of two histidine residues (*Ka*His110 and *Ka*His112), which is not conserved in *Hp*UreE or *Sp*UreE (Colpas *et al*., 1999, Song *et al*., 2001).

Given the important role that UreE has in urease activation and subsequent pathogenicity, we sought to structurally characterise *P. mirabilis* UreE (*Pm*UreE) and investigate its metal binding capacity. Here we present a 2.0 Å crystal structure of *Pm*UreE and show that it can bind five nickel ions per dimer, with the histidine-rich C-terminal tails contributing to the coordination of two Ni^2+^ per dimer. This work increases our knowledge about the diversity of UreE homologues and provides insight into how UreE binds nickel within *P. mirabilis*.

## 2. Materials and methods

### 2.1. Sequence alignment

A multiple-sequence alignment of UreE from *P. mirabilis* (Uniprot ID: P17090), *K. aerogenes* (Uniprot ID: P18317), *H. pylori* (Uniprot ID: B6JPH3), and *S. pasteurii* (Uniprot ID: P50049) was performed using ClustalW (Higgins *et al*., 1996) and visualised with ESPript3 (Gouet *et al*., 1999).

### 2.2. Molecular cloning and mutagenesis

To generate the *Pm*UreE expression plasmid (pETHis6TEVLIC_UreE), the *P. mirabilis* ATCC 12453 *ureE* open reading frame (486 bp) and the kanamycin-resistant pETHis6TEVLIC plasmid backbone were amplified separately by PCR using Q5^®^ High-Fidelity 2× Master Mix (New England Biolabs) (primers listed in Table S1). The pETHis6TEVLIC cloning vector (1B) was a gift from Scott Gradia (Addgene plasmid # 29653; http://n2t.net/addgene:29653; RRID: Addgene_29653). Reaction setup and reagent volumes followed the vendor’s instructions. In the amplification of the backbone, the region encoding the N-terminal His6-tag and TEV protease cleavage site was removed. The insert and backbone were assembled using NEBuilder^®^ HiFi DNA Assembly Master Mix. Chemically competent *E. coli* Top10 cells were transformed with the assembly. Transformants were selected on kanamycin plates, and colonies were screened by PCR with NEBuilder Quick-Load^®^ Taq 2× Master Mix.

To generate the *Pm*UreEΔC17 truncation mutant expression plasmid, a stop codon was inserted into the pETHis6TEVLIC_UreE construct at the intended truncation site (after Gly144), utilising the New England Biolabs site-directed mutagenesis protocol, with the Q5^®^ High-Fidelity 2× Master Mix (primers in Table S1) and KLD Enzyme Mix. The resulting product was transformed into chemically competent NEB *E.coli* Turbo cells and plated on kanamycin plates.

For both expression constructs, plasmid DNA of positive clones was isolated using the NEB Monarch^®^ Spin Plasmid Miniprep Kit. The plasmid sequences were confirmed by whole-plasmid Nanopore sequencing (Plasmidsaurus) (Fig. S2).

### 2.3. Protein expression and purification

*E. coli* BL21(DE3) pLysS transformed with the pETHis6TEVLIC_UreE plasmid or *E. coli* BL21 (DE3) carrying pETHis6TEVLIC_UreEΔC17 was cultured overnight in LB media supplemented with 50 µg/mL kanamycin and 34 µg/mL chloramphenicol (for the cells with the pLysS plasmid). The starter culture was diluted 1:100 into 1 L Terrific Broth (TB) containing the same antibiotic concentrations. Cultures were incubated at 310 K and 180 rpm until the optical density at 600 nm (OD_600_) reached 0.8-1.0. Protein expression was induced with 1 mM IPTG (isopropyl β-D-1-thiogalactopyranoside), followed by further incubation at 310 K for 4 h. Cells were harvested by centrifugation at 4,500 rpm (Beckman Coulter Avanti JE, JLA-9.1000 Rotor) for 15 min at 277 K. Both proteins were expressed without affinity tags and were purified following a similar method reported previously (Sriwanthana *et al*., 1994).

All purification steps were performed at 277 K unless otherwise specified. Cell pellets were resuspended in Buffer A (20 mM Tris-HCl, pH 7.5; 500 mM NaCl) supplemented with a protease-inhibitor tablet (SIGMAFAST™ Protease Inhibitor Tablets, Sigma-Aldrich) and DNase I, and lysed using a Qsonica Q700 probe sonicator (cycles of 10 s on/20 s off, total 5 min, 40% amplitude). The lysate was clarified by centrifugation (30,000 × g, 30 min), and the supernatant was loaded onto 2 or 4 mL of Ni-NTA His-Bind^®^ Resin (EMD Millipore; cat. 70666-5) pre-equilibrated in Buffer A and packed in a gravity-flow column. After binding, the column was washed with Buffer B (Buffer A + 20 or 60 mM imidazole), and bound protein was eluted with Buffer C (Buffer A + 250 mM imidazole). Eluted fractions were pooled and concentrated in a 3 kDa MWCO centrifugal concentrator, filtered (0.22 µm), and subjected to size-exclusion chromatography on a HiLoad 16/600 Superdex 75 column (ÄKTA Pure system, Cytiva) pre-equilibrated in SEC buffer (20 mM HEPES, pH 7.5; 100 mM NaCl, with or without 1 mM EDTA). Elution was monitored at 280 nm, and peak fractions were analysed by SDS-PAGE on Bolt™ 4-12% Bis-Tris Plus Protein Gels in MES buffer (Thermo Fisher Scientific). Fractions corresponding to dimer *Pm*UreE/*Pm*UreE-ΔC17 were collected. The purified protein was concentrated in 3 kDa MWCO concentrators, and then aliquoted, flash-frozen in liquid nitrogen, and stored at 193 K. Total yield for *Pm*UreE was ∼109 mg from 4.5 g of cell pellet and *Pm*UreE-ΔC17 was ∼41 mg from 3.4 g of cell pellet.

*Pm*UreE *and Pm*UreE-ΔC17 concentrations were determined by measuring the absorbance at 280 nm using a QuickDrop spectrophotometer (SpectraMax) and dividing the measurement by the ProtParam generated theoretical extinction coefficient of 0.736 or 0.728, respectively.

### 2.4. Crystallisation

The *Pm*UreE protein used for crystallisation was purified without the addition of EDTA. Initial crystallisation trials were performed by vapour-diffusion sitting drops on 96-well MRC 2-well plates using a Formulatrix NT8 microfluidics protein crystallisation robot. Four commercial screens were tested across protein concentrations ranging from 3 to 20 mg/mL: Shotgun (Molecular Dimensions)(Abrahams & Newman, 2021), JCSG-plus (Molecular Dimensions), PEGRx (Hampton Research), and Crystal Screen HT (Hampton Research). Drops were set at 100 nL protein + 100 nL reservoir solution (1:1), equilibrated against 50 µL reservoir solution and incubated at 293 K. Needle-shaped crystals were obtained in 0.2 M ammonium citrate dibasic, 20% w/v polyethylene glycol (PEG) 3350 (JCSG-plus condition A3) with 5 mg/ml *Pm*UreE within 5-7 days. No other crystallisation hits were obtained in the screens. Optimisation trials around the initial hit were conducted using 24-well VDX plates with sealant (Hampton Research) by hanging drop vapour-diffusion at 293 K with drops containing 1 µL protein and 1 µL or 2 µL reservoir solution equilibrated against 500 µL reservoir solution. As systematic variation of PEG percentage and salt concentration and additives produced only modest improvements, micro-seeding was performed as reported previously (D’Arcy *et al*., 2014, Bergfors, 2007). The seed stock was prepared from needle-like crystals grown in 0.14-0.22 M Ammonium citrate dibasic; 16-22 % w/v PEG 3350 by crushing the crystals in a small volume of mother liquor, this was then serially diluted 100, 1,000, and 10,000-fold.

Robot-assisted re-screening was then performed across the same four screens (Shotgun, JCSG-plus, PEGRx, Crystal Screen) at a protein concentration of 5 mg/mL using a sitting-drop setup of 100 nL protein, 100 nL reservoir solution, and 50 nL seed stock at 293 K. Both undiluted seed stock and a 100-fold dilution were tested. Many more hits were obtained and those conditions that produced well-formed crystals were then transferred to a 24-well hanging-drop optimisation tray with seeding, using 1 µL protein, 1 µL reservoir solution, and 0.50 µL seed stock or 1 µL + 1 µL + 0.25 µL. During 24-well plate optimisation, 100-fold seed caused over-nucleation (dense microcrystals), so 1,000- and 10,000-fold dilutions were used for subsequent drops. During optimisation, 5% glycerol was added as an additive in some trials to stabilise the crystals. In parallel, the published crystallisation conditions reported for UreE homologues were also tested as additional leads (Shi *et al*., 2010). Large rod-shaped crystals were obtained after 1-3 days in 0.1 M sodium citrate pH range 5.0-5.5, 13-19% w/v PEG 8000 or PEG 6000, with or without 5% w/v glycerol and were selected for X-ray diffraction analysis. Despite the morphological improvements, the crystals were fragile, so to limit handling, they were cryoprotected by adding 2 µL of 40% (v/v) glycerol directly to the crystallisation drop. Crystals were mounted and immediately flash-frozen in liquid nitrogen before being stored at cryogenic temperatures. The best diffracting (1.86 Å) crystal was grown in 0.1 M sodium citrate pH 5, 15% PEG 6000, 5 % glycerol.

### 2.5. X-ray data collection, structure solution and refinement

X-ray diffraction data were collected at the MX2 microfocus macromolecular crystallography beamline at the Australian Synchrotron (Melbourne, Victoria, Australia) (McPhillips *et al*., 2002) through remote access using the HTML5 Virtual Network Computing (VNC) client, Guacamole. 3600 frames of 0.1° rotation with a total exposure time of 36 s were collected using an Dectris Eiger 16M detector at a wavelength of 0.9537 Å. Raw data were automatically processed at the beamline using XDS (X-ray Detector Software) (Kabsch, 2010) in the space group *P*2_1_2_1_2_1_ and used as input for re-scaling in *AIMLESS* (Evans & Murshudov, 2013), as implemented in the Collaborative Computational Project No. 4 (CCP4) suite (version 2.8.5)(Agirre *et al*., 2023). The high-resolution cutoff was set to 2.0 Å to improve *I/σ(I)* and R statistics. Expected solvent content and the likely number of macromolecular copies per asymmetric unit were estimated with the Matthews coefficient tool in CCP4 (Matthews, 1968, Kantardjieff & Rupp, 2003).

Molecular replacement (MR) was carried out with Phaser-MR (maximum-likelihood MR) (McCoy *et al*., 2007) in the PHENIX suite (Python-based Hierarchical ENvironment for Integrated Xtallography; version 1.21.2-5419) (Liebschner *et al*., 2019, Adams *et al*., 2010). The search model was the AlphaFold prediction of *Pm*UreE from the UniProt database (AF-P17090-F1, Uniprot ID: P17090) (Jumper *et al*., 2021). The model was prepared in Phenix *Process Predicted Model* to remove low confidence regions and replace values in the B-factor field. Candidate copy numbers suggested by the Matthews analysis were tested, and the solution was selected based on translation-function Z-score (TFZ) and log-likelihood gain (LLG).

Initial model refinement was performed in PHENIX.refine (Afonine *et al*., 2012). Manual model building based on the refined structure was performed in Coot (Crystallographic Object-Oriented Toolkit; version 0.9.8.95) (Emsley *et al*., 2010). Side-chain conformations, backbone geometry, and alternate conformations were adjusted against the electron density maps. Each manual editing cycle was followed by an additional round of refinement in PHENIX until no significant features remained in the difference maps and the model statistics converged. Further refinement included adding water molecules where the difference in density and hydrogen-bonding geometry supported placement. Model validation was performed using MolProbity (Williams *et al*., 2018) (as implemented in PHENIX) to assess Ramachandran statistics, rotamers, clashscore, and overall geometry. Data-to-model agreement was monitored throughout using R_work_/R_free_. Coordinates and structure factors were deposited to the Protein Data Bank with the accession ID 9ZLO and crystallographic statistics are shown in Table 1. Structural alignments were performed in Coot using the ‘Secondary Structure Matching (SSM) Superpose’ command for homologous structures and ‘Least Squares (LSQ) Superpose’ for *Pm*UreE chains (Emsley *et al*., 2010). Figures were made using PyMOL (Schrödinger, LLC; version 3.1.6.1).

**Table 1.**
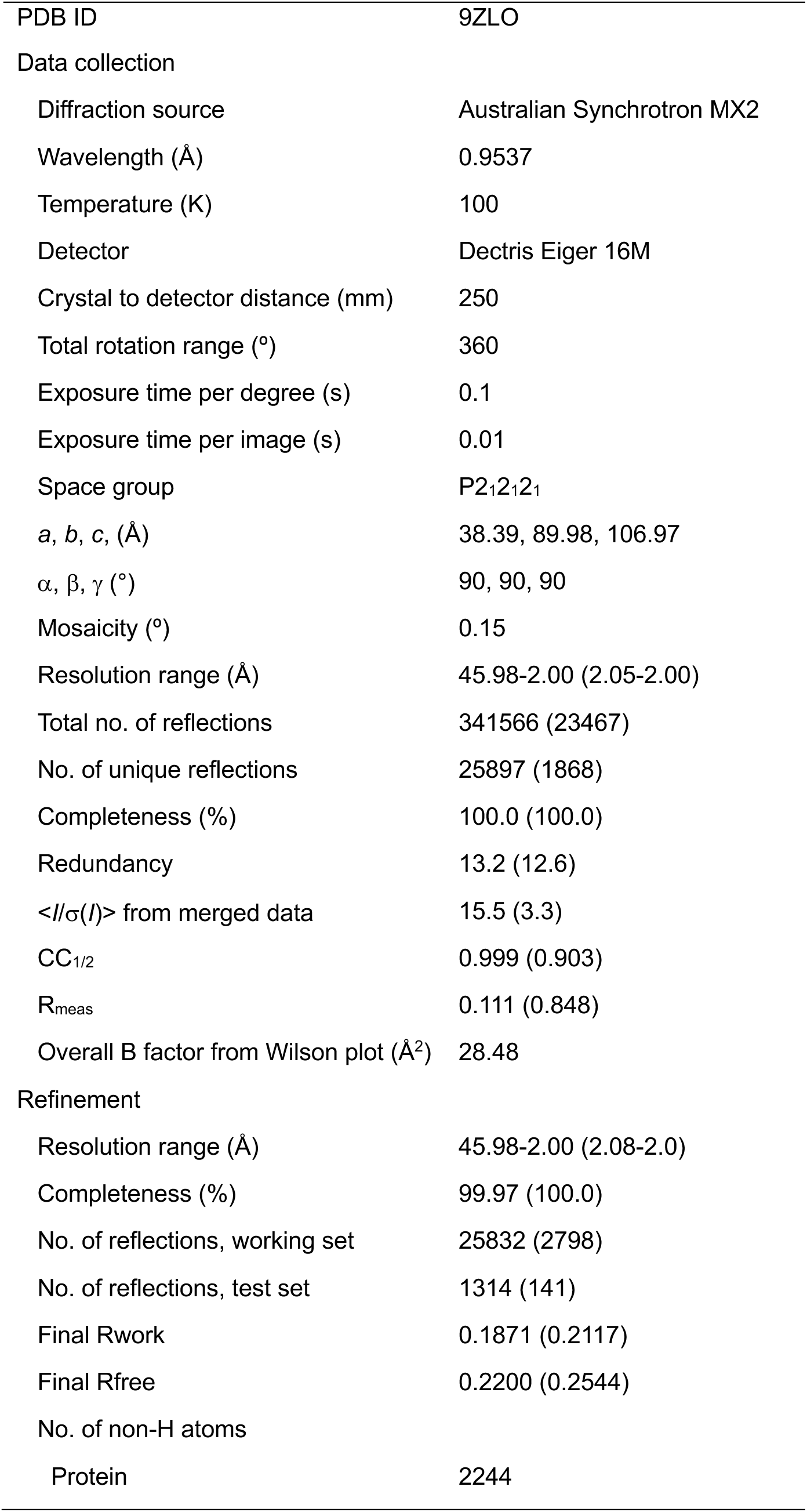
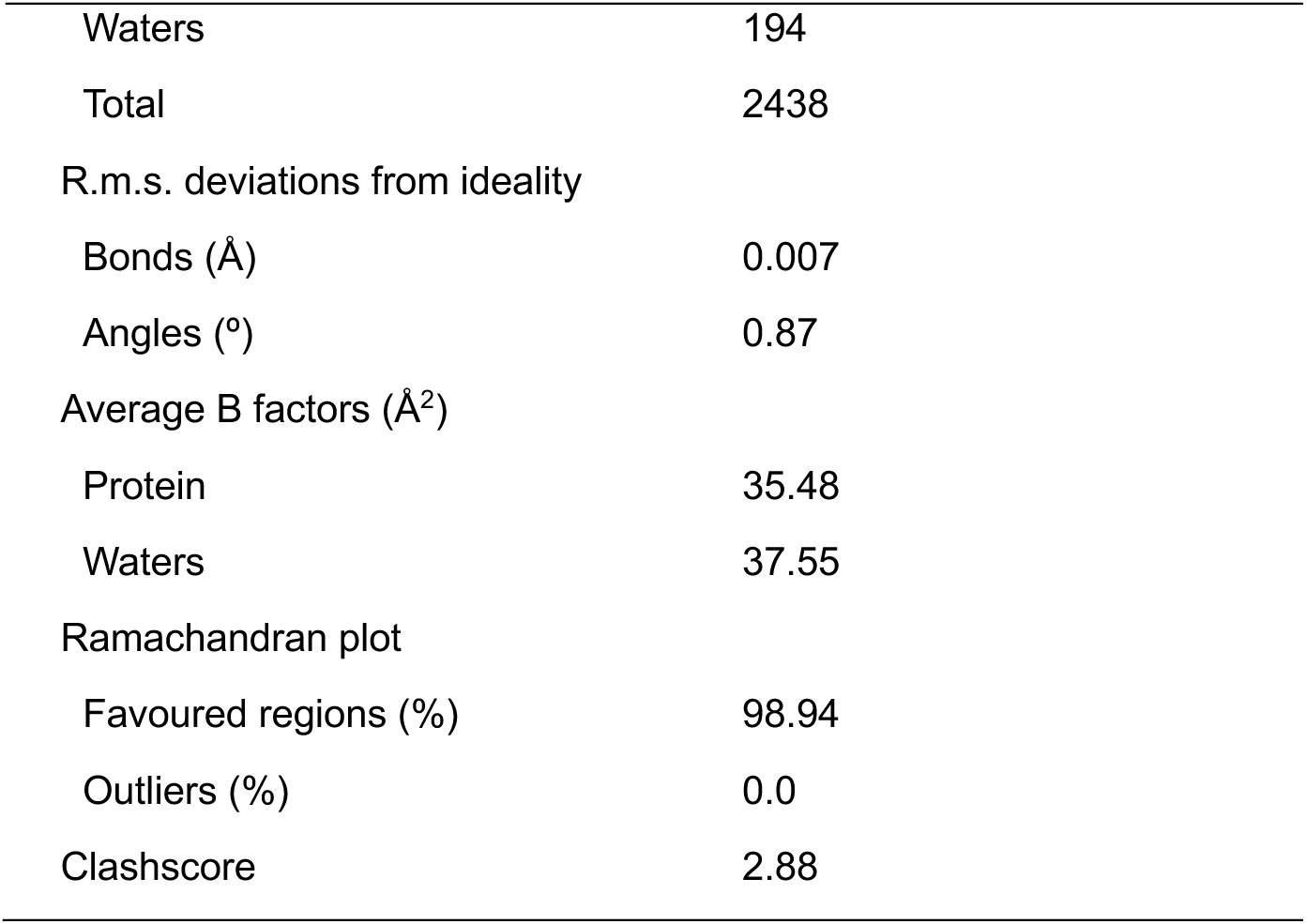
Crystallographic statistics for the PmUreE structure Values in parentheses are for the highest resolution shell.

|  |  |
| --- | --- |
| PDB ID | 9ZLO |
| Data collection |  |
| Diffraction source | Australian Synchrotron MX2 |
| Wavelength (Å) | 0.9537 |
| Temperature (K) | 100 |
| Detector | Dectris Eiger 16M |
| Crystal to detector distance (mm) | 250 |
| Total rotation range (°) | 360 |
| Exposure time per degree (s) | 0.1 |
| Exposure time per image (s) | 0.01 |
| Space group | P2<sub>1</sub>2<sub>1</sub>2<sub>1</sub> |
| *a*, *b*, *c*, (Å) | 38.39, 89.98, 106.97 |
| $\alpha$, $\beta$, $\gamma$ (°) | 90, 90, 90 |
| Mosaicity (°) | 0.15 |
| Resolution range (Å) | 45.98-2.00 (2.05-2.00) |
| Total no. of reflections | 341566 (23467) |
| No. of unique reflections | 25897 (1868) |
| Completeness (%) | 100.0 (100.0) |
| Redundancy | 13.2 (12.6) |
| $\langle I/\sigma(I) \rangle$ from merged data | 15.5 (3.3) |
| CC<sub>1/2</sub> | 0.999 (0.903) |
| R<sub>meas</sub> | 0.111 (0.848) |
| Overall B factor from Wilson plot (Å<sup>2</sup>) | 28.48 |
| Refinement |  |
| Resolution range (Å) | 45.98-2.00 (2.08-2.0) |
| Completeness (%) | 99.97 (100.0) |
| No. of reflections, working set | 25832 (2798) |
| No. of reflections, test set | 1314 (141) |
| Final Rwork | 0.1871 (0.2117) |
| Final Rfree | 0.2200 (0.2544) |
| No. of non-H atoms |  |
| Protein | 2244 |
| Waters | 194 |
| Total | 2438 |
| R.m.s. deviations from ideality |  |
| Bonds (Å) | 0.007 |
| Angles (°) | 0.87 |
| Average B factors (Å<sup>2</sup>) |  |
| Protein | 35.48 |
| Waters | 37.55 |
| Ramachandran plot |  |
| Favoured regions (%) | 98.94 |
| Outliers (%) | 0.0 |
| Clashscore | 2.88 |

### 2.6. Inductively Coupled Plasma Mass Spectrometry (ICP-MS) quantification of nickel binding

Quantification of nickel ions associated with *Pm*UreE and *Pm*UreE-ΔC17 was determined using ICP-MS. Samples (typically 20 µM protein in 500 µL incubation buffer) were incubated with 100 µM NiCl_2_ at room temperature for 2.5 h, then dialysed overnight at 277 K against 1 L of dialysis buffer to remove free metal ions. Protein concentration was measured post-dialysis for subsequent stoichiometry calculations. For digestion, ∼10 µM protein was mixed 1:1 with 4% (v/v) nitric acid (HNO_3_) and incubated overnight at room temperature to release bound metal. Process blanks (dialysis buffer and 4% HNO_3_) were prepared in parallel to test background metal content. Digests were centrifuged (20,000 × g, 10 min, room temperature), and supernatants were collected for analysis.

The analysis was performed on a Thermo Scientific iCAP RQ ICP-MS system in KEDS (Kinetic Energy Dispersion Sensitive) mode using helium as a collision gas. A multi-element standard was used for ^60^Ni calibration (Agilent IntelliQuant 68 multi-element standard, “IQ-1”, #5190-9422). A five-point external calibration was prepared for ^60^Ni at concentrations of 1, 10, 50, 100, and 500 ppb (µg L-1) in the same acid matrix as the samples (2% HNO_3_), yielding a linear regression coefficient (R^2^) of ≥ 0.999. The software Qtegra was used to process the data. A mid-range quality control (QC) standard was prepared independently from the calibration standards and measured before and after the sample sequence to assess and correct for instrumental drift during run time. Instrumental drift correction was performed using linear interpolation where the rate of drift was calculated from the rate of change in 100 ppb QC values obtained at the beginning and the end of the run queue and was applied to each sample as a function of its position in the queue. Each sample was analysed in triplicate and the average used for calculations. Limit of detection (LOD) and limit of quantification (LOQ) were defined from measured blanks (in ppb) as: LOD = 3 × Standard Deviation (SD)_blank, LOQ = 10 × SD_blank (SD_blank from ≥ 3 independent blank digests). Measurements below LOD were flagged as Below Detection Limit (BDL) and excluded from stoichiometry calculations.

Instrument output (µg L^-1^, ppb) was corrected for the total dilution factors to obtain the ^60^Ni concentration in the post-dialysis sample. Where required, concentrations were converted to micromolar using the molar mass of ^60^Ni, and nickel-to-protein ratios were calculated using the measured protein concentration and the oligomeric state specified for the experiment (see formula below). Results are reported as mean ± SD of three independent experiments.

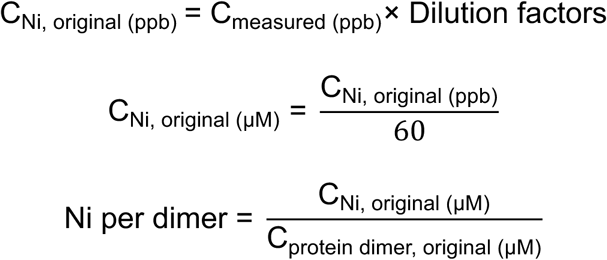

## 3. Results and discussion

### 3.1. *Pm*UreE crystallisation and structure analysis

The full length *Pm*UreE protein was purified by immobilized metal affinity chromatography using its native ability to bind nickel, followed by size-exclusion chromatography (SEC) (Fig. S3). The elution volume on SEC approximately corresponded to the molecular weight of a homodimeric species, in line with previous purifications of *Pm*UreE and other UreE homologues (Heimer & Mobley, 2001, Sriwanthana *et al*., 1994). Despite the long, unstructured, histidine-rich C-terminus of the full length UreE, crystals formed in the initial sparse matrix screens and were optimised to large, rod-like morphologies using micro-seeding (Fig. S4). The best crystal diffracted to 1.86 Å, though the data processing statistics were improved by excluding diffraction data greater than 2.0 Å. The structure was solved using molecular replacement with the AlphaFold model for *Pm*UreE (AF-P17090-F1) and two copies of the molecule were found in the asymmetric unit (RMSD 0.8 Å, 141 Cα atoms aligned). Each of these chains formed the biological dimer through crystallographic symmetry (Fig. 1). In both chains, continuous density is observed for the folded cores, whereas flexible regions near the termini are not resolved: the C-terminal tails of both chains are absent from the maps, and Chain B lacks clear density for residues 19-21 near the N-terminus (Fig. S5*a*). The final models extend to residue 147 in Chain A and to residue 144 in Chain B. Each monomer adopts a two-domain structure connected by a short linker (Fig. 1*a*). The N-terminal domain (NTD) consists of two short α-helices and two three-stranded mixed β-sheets that stack almost orthogonally to form a rigid core. The C-terminal domain (CTD) comprises an antiparallel four-stranded β-sheet flanked by two α-helices (Fig. 1*a*). The biological dimer is formed by a head-to-head association of the CTD α3-helices (Pro89-Arg102) (Fig. 1*b*,*c*), with the dimerization stabilised by a combination of hydrophobic and hydrogen-bonding interactions between α3 and the β8 (His103-Ala110) of the neighbouring protomer (Fig. 1*d*). The resolved region of the flexible C-terminus also interacts with the α3 of the adjacent protomer (Fig. 1*d*).

**Figure 1.**
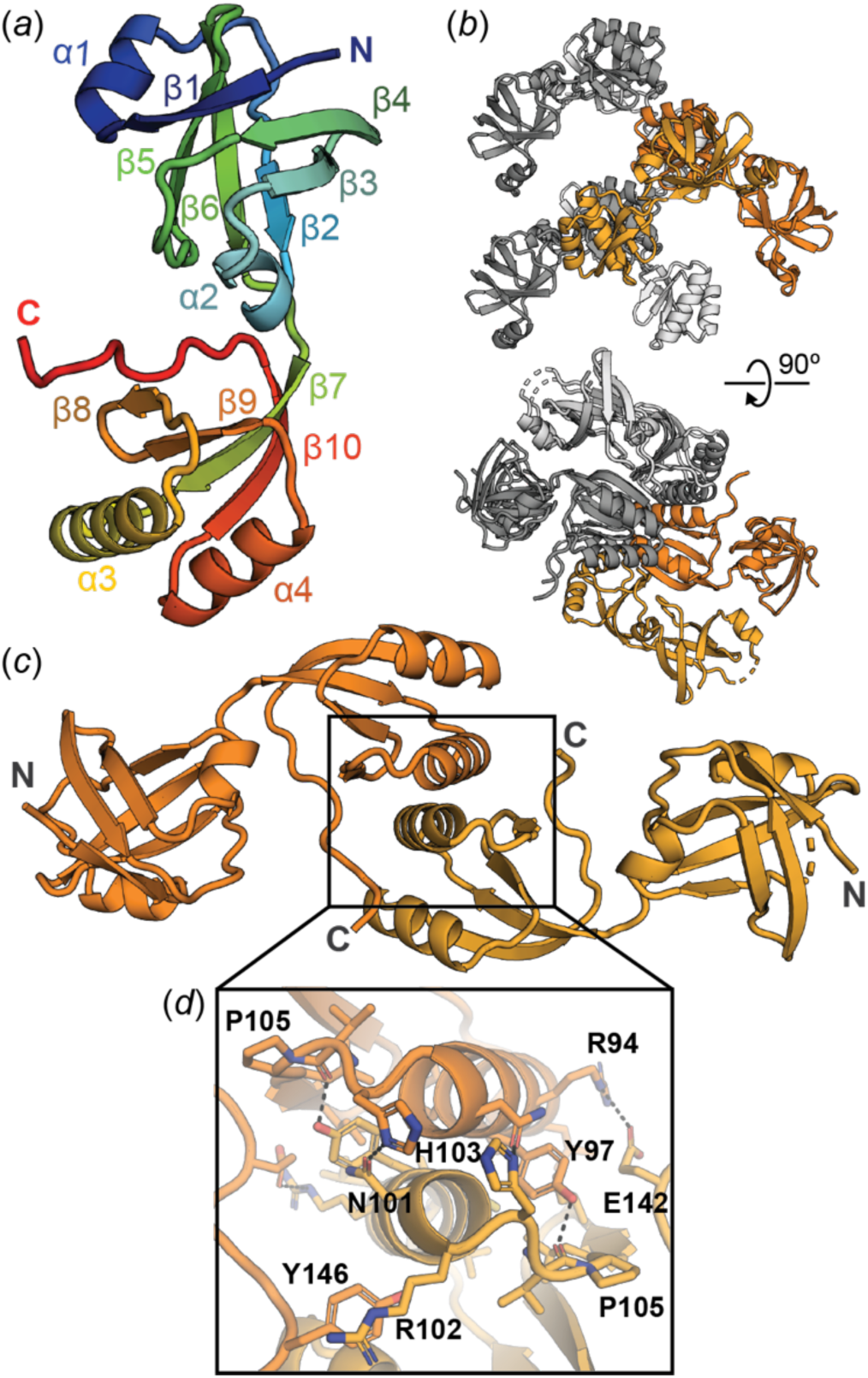
Crystal structure of *Pm*UreE. (*a*) Cartoon representation of *Pm*UreE with secondary structure features labelled and coloured in rainbow, with the N-terminal region shown in blue/green and C-terminal region shown in orange/red. (*b*) Crystal packing of the *Pm*UreE crystal structure. The asymmetric unit is coloured in orange (chain A is darker orange; chain B is lighter orange) while symmetry mates are coloured in grey (chain A equivalents are dark grey; chain B are light grey). The biologically relevant dimer is formed through crystallographic symmetry. (*c*) Cartoon representation of the *Pm*UreE dimer with chain A coloured dark orange and chain B coloured light orange. (*d*) Closer view of the dimerization interface, with key residues labelled and selected hydrogen bonding interactions represented with dashed black lines.

Inspection of the electron-density maps did not reveal convincing density for bound metal ions. No strong, spherical density consistent with a transition metal was observed at the dimer interface or elsewhere, and coordination geometry consistent with Ni^2+^ was not evident. The crystal structure, therefore, represents an apo form of *Pm*UreE under the conditions used for crystallisation. Instead, two irregular, relatively strong density blobs were observed near the N-terminal region of the monomers. Their shapes were elongated/ill-defined rather than spherical. Given the crystallisation solution, these features are most consistent with bound buffer components (e.g., citrate). However, attempts to model them and refine candidate ligands or ions in Phenix were not successful. As a result, these sites were left unmodelled in the final coordinates.

### 3.2. *Pm*UreE shares structural similarity with other UreE homologues

To compare the *Pm*UreE structure to other solved UreE structures, superpositions were generated against *K. aerogenes* UreE (*Ka*UreE; PDB: 1GMU), *H. pylori* UreE (*Hp*UreE; PDB: 3L9Z), and *S. pasteurii* UreE (*Sp*UreE; PDB: 1EAR) (Fig. 2*a*). *Pm*UreE shares the greatest structural similarity with *Ka*UreE (RMSD 1.1 Å, 134 Cα atoms aligned), but all four proteins share a conserved two-domain structure and a head-to-head dimer arrangement. Residues at the dimerization interface (*Pm*Pro89-Ala110) are well conserved between *K. aerogenes* and *P. mirabilis* (Fig. S1). The overlays indicate that the folded cores align well, with deviations mostly in the N-terminal region and a few solvent-exposed loops (Fig. 2*a*). Intriguingly, *Pm*UreE has an additional short but well-defined α-helix (α1, Fig. 1*a*, S5*b*) in the N-terminal domain, that is a flexible loop in other solved structures. Most studies have focused on the metal binding capabilities of UreE rather than the role of the NTD; however, the NTD is structurally related to heat shock proteins and has been referred to as a peptide binding domain (Song *et al*., 2001). A recent study showed that the NTD of *Hp*UreE is involved in binding to *Hp*UreG during nickel transfer between the two proteins (Chan *et al*., 2025). Though the interaction interface between *Hp*UreE and *Hp*UreG is on the opposite side to α1 in *Pm*UreE, it is possible that this helix could be involved in specialised protein-protein interactions for the UreE acquisition or delivery of nickel within *P. mirabilis*.

**Figure 2.**
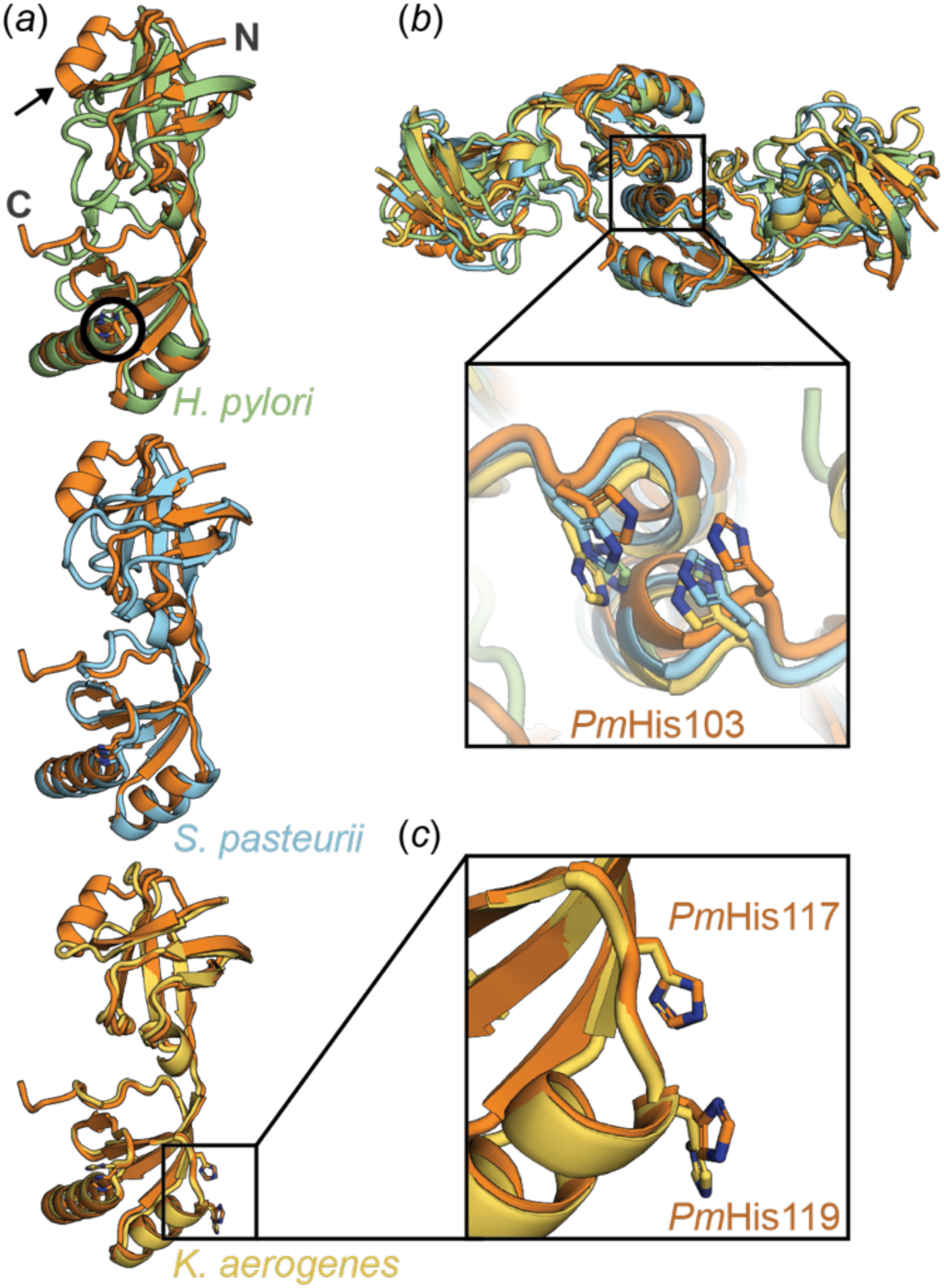
Structural alignment of *Pm*UreE with other UreE homologues. (*a*) Alignment of *H. pylori* UreE (green, PDB: 3L9Z, RMSD 2.1 Å, 118 Cα atoms aligned), *S. pasteurii* UreE (blue, PDB: 1EAR, RMSD 2.1 Å, 120 Cα atoms aligned) and *K. aerogenes* (yellow, PDB: 1GMU, RMSD 1.1 Å, 134 Cα atoms aligned) against *Pm*UreE (orange). Arrow indicates the additional α-helix in *Pm*UreE and the black circle highlights the position of the conserved histidine residue that sits at the dimer interface (*b*) Alignment of the dimeric forms of the UreE homologues (coloured as per (*a*)) with the Ni^2+^ binding site at the dimerization interface featured (inset). (*c*) Intramolecular, secondary Ni^2+^ binding site is conserved between *Pm*UreE and *Ka*UreE.

Given the overall conservation in architecture, we focused our analyses on the two histidine-rich sites that are known to have functional roles in UreE homologues. The first is located at the dimer interface and has been implicated in metal binding across all three homologues. In the *Pm*UreE dimer, *Pm*His103 from each monomer occupies the same interfacial site as the equivalent residues in *Ka*UreE (*Ka*His96), *Hp*UreE (*Hp*His102), and *Sp*UreE (*Sp*His100) (Fig. 2*b*). The side-chain orientations are closely matched across the four structures, forming a conserved Ni^2+^ binding site at the CTD-CTD interface. Mutation experiments have shown that these dimer interface histidine residues are essential for urease activation in *K. aerogenes* and *H. pylori*, indicating that they are involved in the transfer of nickel to the downstream protein, UreG (Colpas *et al*., 1999, Colpas & Hausinger, 2000, Shi *et al*., 2010). This was confirmed by the recently released structure of the *Hp*UreE-*Hp*UreG interaction (Chan *et al*., 2025).

The second histidine-rich site resides within each C-terminal domain and is associated with metal-binding in *K. aerogenes* but is not present in *H. pylori* or *S. pasteurii*. Within each *Pm*UreE monomer, *Pm*His117 and *Pm*His119 align with *Ka*His110 and *Ka*His112 in the CTD (Fig. 2*c*). The relative positions of the four-stranded β-sheet and adjacent helices are conserved, creating an intramolecular histidine pair analogous to the Ni^2+^ binding site reported for *Ka*UreE (Song *et al*., 2001). The *Ka*UreE structure was solved with metal bound in these sites (Song *et al*., 2001) and mutation of *Ka*His110/*Ka*His112 weakens Ni^2+^ binding and reduces urease activation *in vivo*, emphasising the role of the intramolecular site in effective metal trafficking (Colpas *et al*., 1999).

### 3.3. *Pm*UreE and *Pm*UreE-ΔC17 nickel binding capacity

Besides the two Ni^2+^ binding sites discussed in 3.2, the histidine-rich C-terminus, although often unresolved in electron density, has been shown in related systems to increase total Ni^2+^ binding capacity and to modulate effective stoichiometry (Lee *et al*., 1993, Brayman & Hausinger, 1996). To investigate the nickel binding stoichiometry of *Pm*UreE and particularly the contribution of the C-terminus, we generated a C-terminally truncated mutant of *Pm*UreE (*Pm*UreE-ΔC17) (Fig. S3*b*) and analysed the nickel binding capacity of both the wild-type *Pm*UreE and *Pm*UreE-ΔC17. Total nickel content was quantified by ICP-MS after NiCl_2_ incubation and dialysis to remove unbound metal. Across three independent preparations, the Ni^2+^ content corresponded to 5.15 ± 0.12 Ni^2+^ per *Pm*UreE dimer and 2.93 ± 0.08 Ni^2+^ per *Pm*UreE-ΔC17 dimer under the assay conditions (Fig. 3*a*). In two of these repeats, the nickel content of the dialysis buffer and 4% HNO_3_ was analysed and found to be BDL or at trace levels (<0.1 µM), so did not contribute to the nickel detected in the samples (Table S2). These results suggest that the C-terminus of *Pm*UreE, which contains eight histidine residues (16 per dimer), contributes to the binding of 2 Ni^2+^.

**Figure 3.**
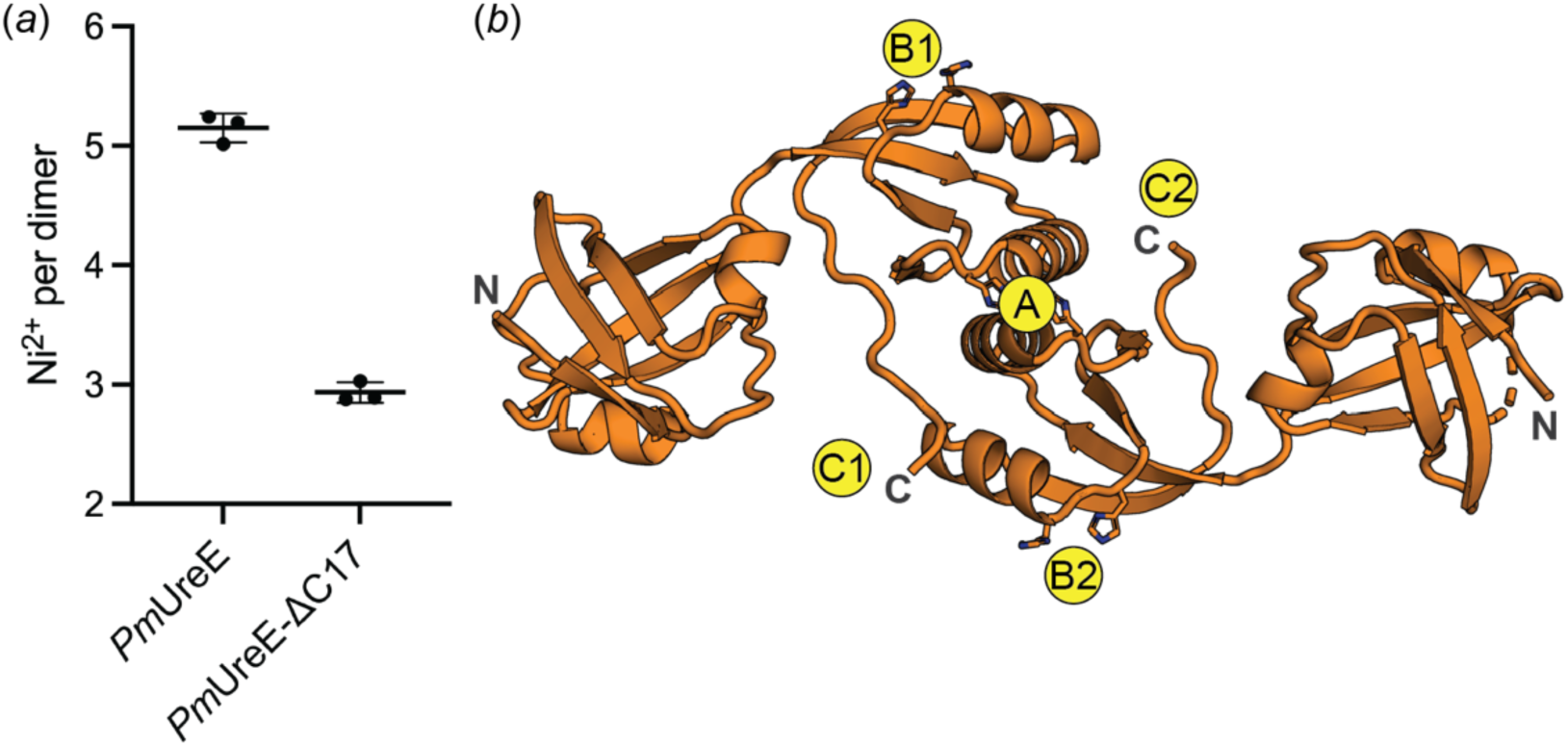
Ni^2+^ binding capacity of *Pm*UreE. (*a*) Quantification of Ni^2+^ binding to *Pm*UreE and *Pm*UreEΛ1C17 using ICP-MS. The mean and standard deviation of the data points from three replicates are shown for each sample. (*b*) Putative Ni^2+^ binding sites in *Pm*UreE based on the ICP-MS results and comparison with previous work on UreE homologues.

Reported stoichiometries by ITC for *Hp*UreE and *Sp*UreE are ∼1 and 2 Ni^2+^ per dimer, respectively (Bellucci *et al*., 2009, Zambelli *et al*., 2013). In line with their lack of a histidine-rich C-terminal tail and the intramolecular CTD binding site, *Hp*UreE and *Sp*UreE show reduced total Ni^2+^ binding compared with *Pm*UreE. As expected from sequence and structural alignment, *Pm*UreE (∼5 Ni^2+^ per dimer by ICP-MS) has the most similar Ni^2+^ binding capacity to *Ka*UreE, which binds ∼6 Ni^2+^ per dimer based on equilibrium dialysis (Lee *et al*., 1993). In *Ka*UreE, the flexible C-terminus has ten histidine residues (contributing 20 per dimer), compared to the eight in *Pm*UreE. A C-terminally truncated form of *Ka*UreE (H144*) binds only ∼2 Ni^2+^ per dimer, which also aligns with our measured data for the *Pm*UreE-ΔC17 mutant (∼3 Ni^2+^ per dimer) (Lee *et al*., 1993, Brayman & Hausinger, 1996). These results suggest that an extra 2 histidine residues in each flexible C-terminus allow the binding of 2 more Ni^2+^ ions per dimer. The C-terminal histidine residues are thought to be involved in maintaining an increased local concentration of Ni^2+^ for transfer to urease. Though the loss of this region results in reduced stoichiometry, removal of the flexible C-terminus has been shown to not affect urease activation in *K. aerogenes* (Mulrooney *et al*., 2005, Brayman & Hausinger, 1996) or the transfer of nickel onto UreG in *H. pylori* (Chan et al., 2025). However, the addition of a histidine-rich C-terminal segment onto *Hp*UreE does increase urease activity and recent work has indicated that the flexible C-terminus is required for the UreE acquisition of nickel from HypA in *H. pylori* (Chan et al., 2025). Whether the C-terminal region has a similar role in other species is yet to be determined.

### 3.4. Putative *Pm*UreE nickel binding sites

The structural alignment of UreE homologues (Fig. 2), along with the measured stoichiometries, supports the assignment of putative Ni^2+^ binding sites in *Pm*UreE (Fig. 3*b*). Site A is assigned to the dimer interface pocket, equivalent to the interface site identified across all UreE structures for metal capture (Remaut *et al*., 2001, Song *et al*., 2001, Shi *et al*., 2010, Banaszak *et al*., 2012, Zambelli *et al*., 2013) (Fig. 3*b*). Sites B1 and B2 correspond to intramolecular sites within each monomer, positioned by the conserved histidine pair that aligns with *Ka*UreE (Fig. 2*c*). Together, the A/B sites account for three core Ni^2+^ binding positions per dimer. The remaining capacity can be explained by the histidine-rich C-terminus: C1 and C2 represent the potential C-terminal-associated Ni^2+^ binding sites, consistent with the high total Ni^2+^ per dimer in the wild type and the loss of ∼2 ions after C-terminal truncation, as observed during ICP-MS (Fig. 3*a*). As there was no corresponding electron density detected for the flexible C-terminus, the precise histidine residues contributing to the C1/C2 sites cannot be assigned. With the eight histidine residues in the *Pm*UreE C-terminal region (Fig. S1), there are several alternative sites or binding modes that could operate, and further experimentation will be required to clarify this.

## 4. Conclusion

Overall, this study has solved the structure of *Pm*UreE at 2.0 Å resolution, identifying a conserved *Pm*UreE dimer scaffold with three conserved sites that likely function in metal binding and transfer. ICP-MS data support multi-site Ni^2+^ binding with a significant contribution from the histidine-rich C-terminal tail, with the ΔC17 truncation decreasing total nickel binding capacity. Together, these data validate *Pm*UreE as a metallochaperone that captures Ni^2+^. We also provide a foundation for understanding the role of UreE in *P. mirabilis* pathogenesis and evaluating its potential as a target for new antibacterial therapies in CAUTIs.

## Acknowledgements

This research was undertaken in part using the MX2 beamline at the Australian Synchrotron, part of Australian Nuclear Science and Technology Organisation, and made use of the Australian Cancer Research Foundation (ACRF) detector. E.J.F.’s laboratory and S.L.M.’s salary was supported by an Australian National University Futures Award. N.T. was supported by a Postgraduate Research Award (ALNSTU25121) from AINSE (Australian Institute of Nuclear Science and Engineering).

## Data availability

The PmUreE crystallographic model and structure factors have been deposited to the Protein Data Bank with the accession code 9ZLO.

## CRediT authorship contribution statement

Jiayi Pan: Writing – original draft, Visualization, Methodology, Investigation, Formal analysis. Sarah L. Mueller: Investigation. Nuren Tasneem: Methodology, Formal analysis. Yang Wu: Methodology, Investigation. Emily J. Furlong: Writing – original draft, Visualization, Funding acquisition, Formal analysis, Conceptualization. All authors contributed to Writing – review & editing.

## Supplementary information

**Table S1.**
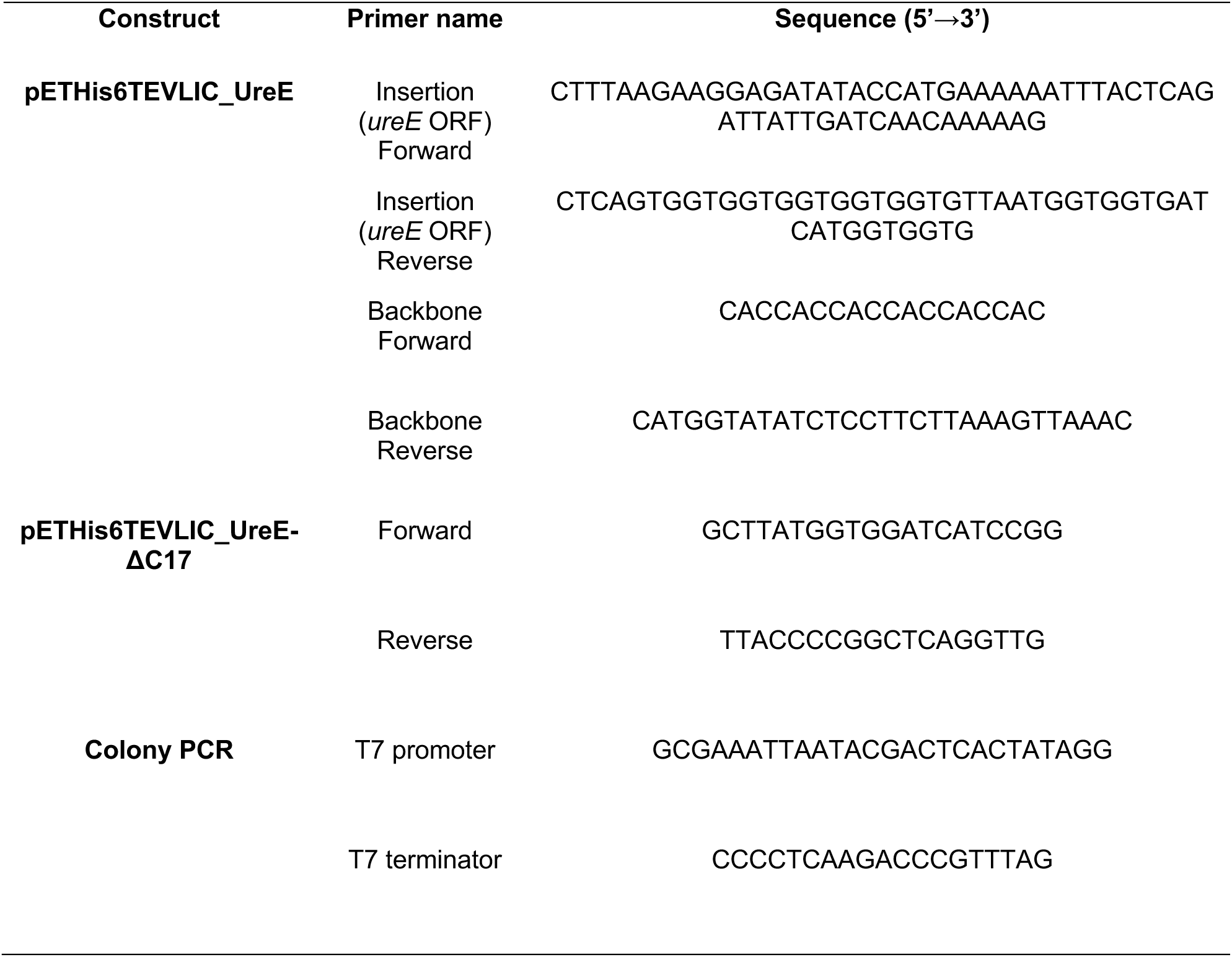
Primers used for cloning. Sequences are shown 5’→3’. “Insertion” primers amplify the gene insert; “Backbone” primers amplify the vector. The ΔC17 reverse primer encodes a stop codon at the truncation site to remove the last 17 amino acid residues. “Forward/Reverse” indicate primer orientation relative to the coding sequence.

**Table S2.** ICP-MS data analysis.

| Sample | Dimeric Protein Conc. (μM) |  |  | Measured Ni<sup>2+</sup> (μM) |  |  | Ni<sup>2+</sup> normalized to 5 μM protein (μM) |  |  | Ni<sup>2+</sup> per dimer |  |  |
| --- | --- | --- | --- | --- | --- | --- | --- | --- | --- | --- | --- | --- |
|  | R1 | R2 | R3 | R1 | R2 | R3 | R1 | R2 | R3 | R1 | R2 | R3 |
| Ni<sup>2+</sup>-incubated protein | 5.49 | 4.36 | 5.94 | 27.53 | 22.63 | 31.14 | 25.07 | 25.95 | 26.2 | 5.014 | 5.19 | 5.24 |
| Dialysis buffer |  | / |  | 0.0658 | NM** | 0.0045 |  | / |  |  | / |  |
| 4 % HNO3 |  | / |  | BDL* | NM** | BDL* |  | / |  |  | / |  |

***PmUreE-ΔC17***
| Sample | Dimeric Protein Conc. (μM) |  |  | Measured Ni<sup>2+</sup> (μM) |  |  | Ni<sup>2+</sup> normalized to 5 μM protein (μM) |  |  | Ni<sup>2+</sup> per dimer |  |  |
| --- | --- | --- | --- | --- | --- | --- | --- | --- | --- | --- | --- | --- |
|  | R1 | R2 | R3 | R1 | R2 | R3 | R1 | R2 | R3 | R1 | R2 | R3 |
| Ni<sup>2+</sup>-incubated protein | 4.47 | 4.18 | 4.77 | 12.86 | 12.65 | 13.78 | 14.38 | 15.13 | 14.43 | 2.88 | 3.03 | 2.89 |
| Dialysis buffer |  | / |  | 0.0022 | NM** | 0.0045 |  | / |  |  | / |  |
| 4 % HNO3 |  | / |  | 0.0038 | NM** | BDL* |  | / |  |  | / |  |

\*Below Detection Limit
\*\* Not Measured

**Figure S1.**
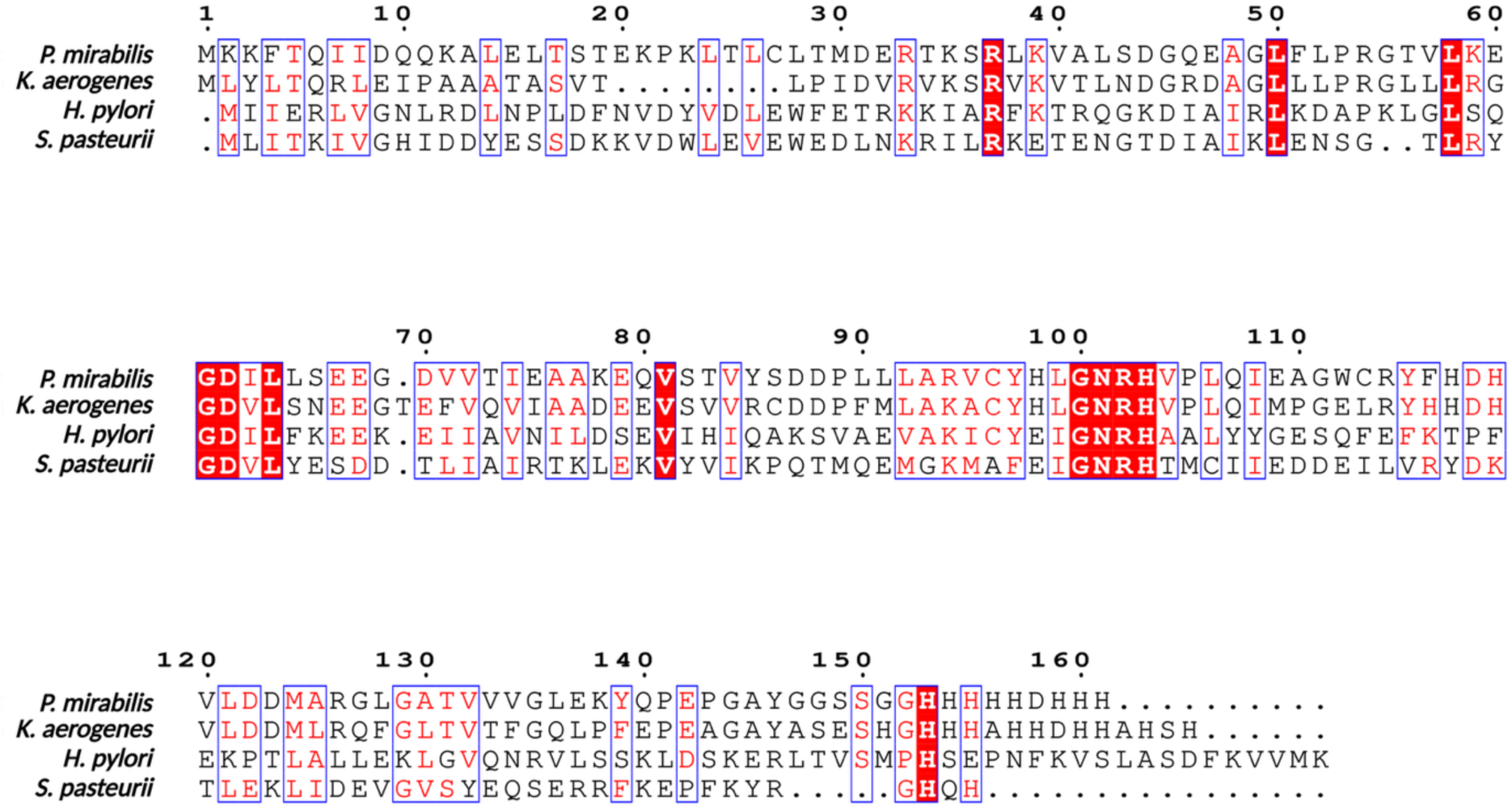
Sequence alignment of UreE homologues. Multiple-sequence alignment of UreE from *P. mirabilis*, *K. aerogenes*, *H. pylori*, and *S. pasteurii*. Residues identical in all four species are highlighted with a red background; residues conserved in three species are outlined with a blue box. Residue numbers follow the *P. mirabilis* sequence.

**Figure S2.**
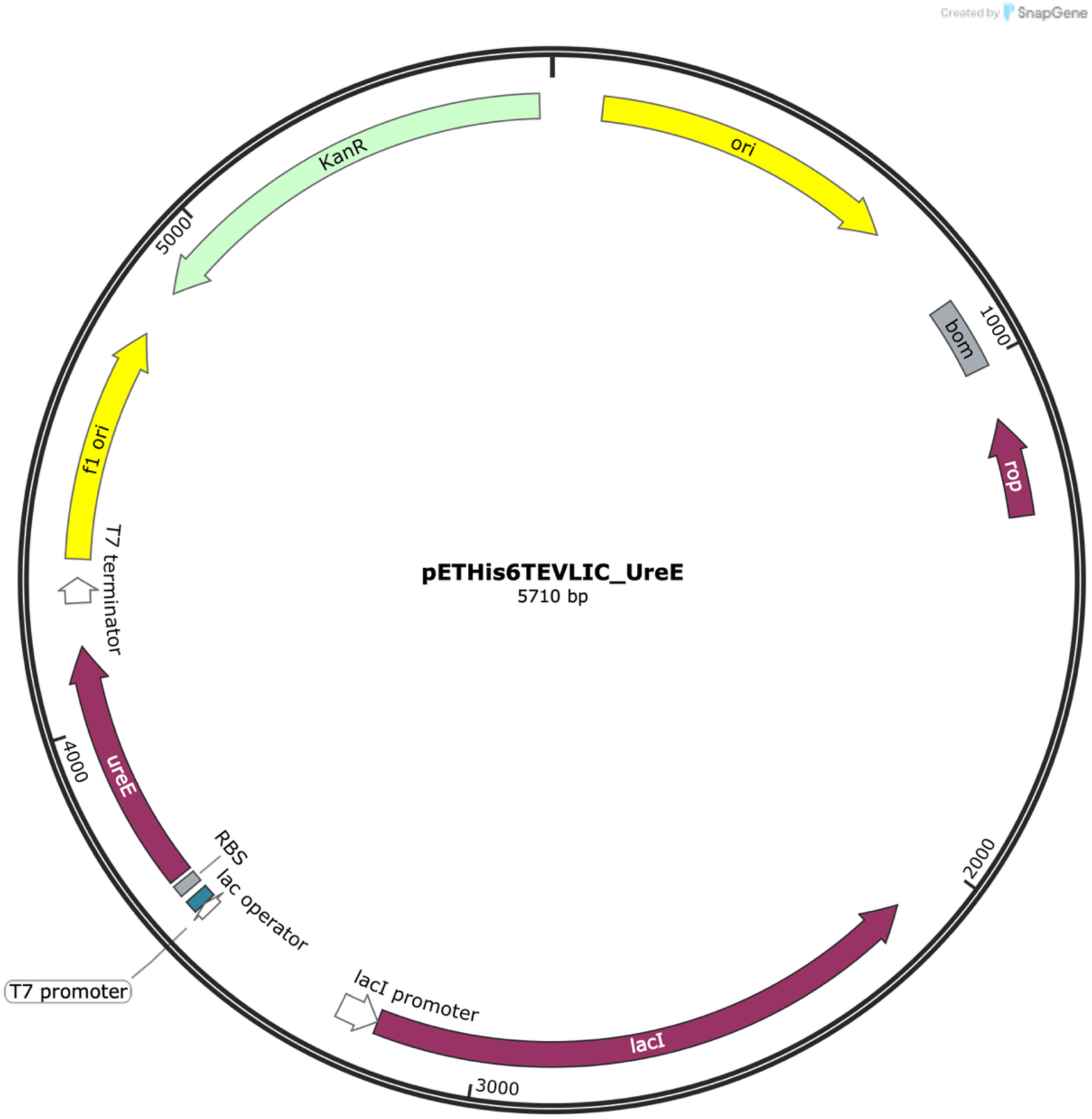
Map of *Pm*UreE expression plasmid. The *Pm*UreE-ΔC17 expression plasmid was identical except for the addition of a stop codon in the DNA after Gly144 in the C-terminal region. Figure created in SnapGene.

**Figure S3.**
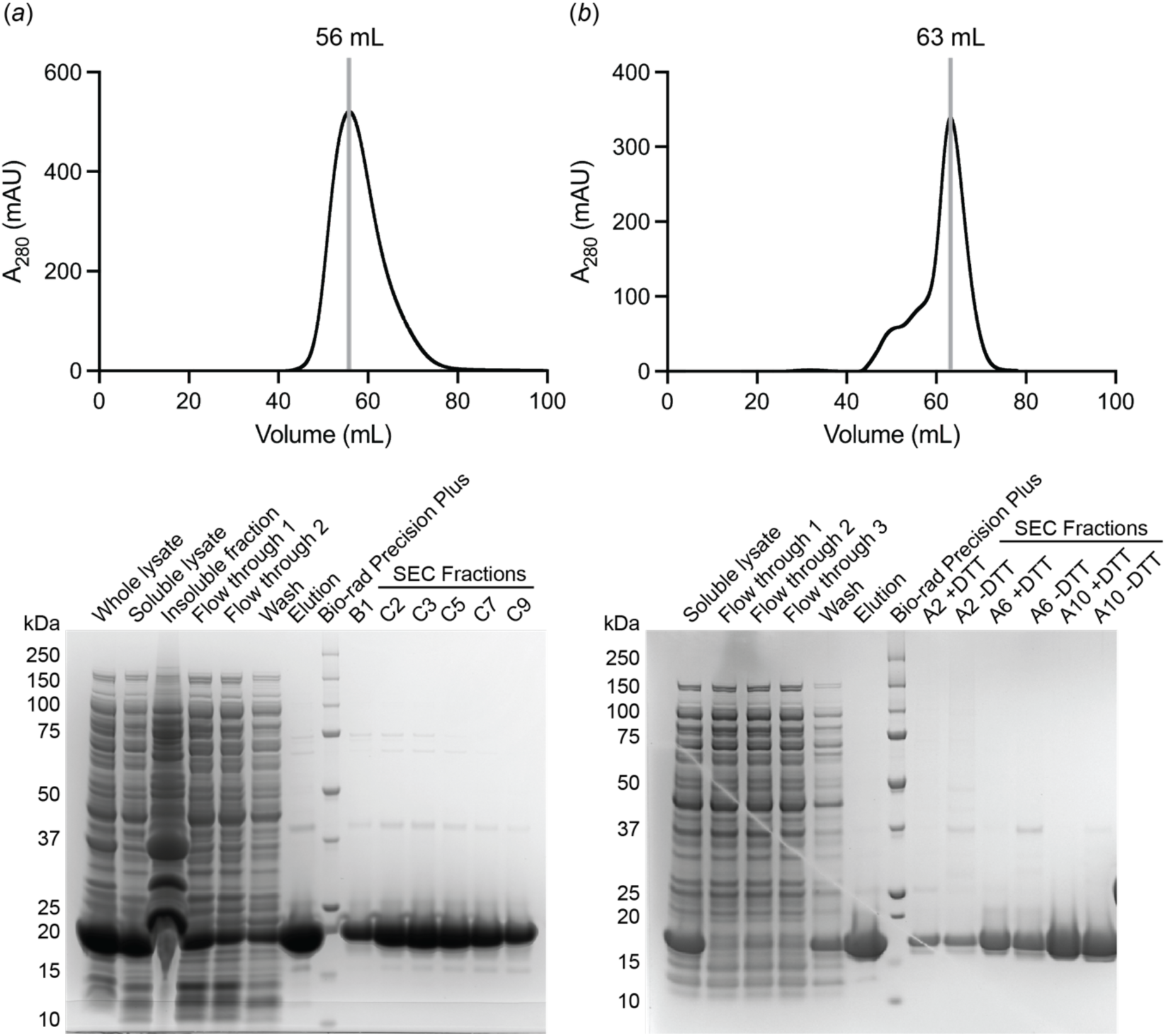
Purification of *Pm*UreE and *Pm*UreE-ΔC17. (*a*) Size-exclusion chromatography (SEC) profile of wild-type *Pm*UreE on a HiLoad 16/600 Superdex 75 column; UV absorbance was monitored at 280 nm. The vertical line marks the elution peak of the protein. Below, SDS-PAGE of all purification steps. SEC fractions showed a single dominant band at ∼18 kDa, corresponding to the expected molecular weight. (*b*) SEC profile of the C-terminal truncation variant *Pm*UreE-ΔC17 on a new HiLoad 16/600 Superdex 75 column with the SDS-PAGE of all purification steps below. SDS-PAGE of the SEC fractions showing a single dominant band at ∼16 kDa, corresponding to the expected monomer molecular weight. The larger than expected difference in elution volume between the proteins is likely due to a new column being used for (*b*), which resulted in better resolution of the peaks.

**Figure S4.**
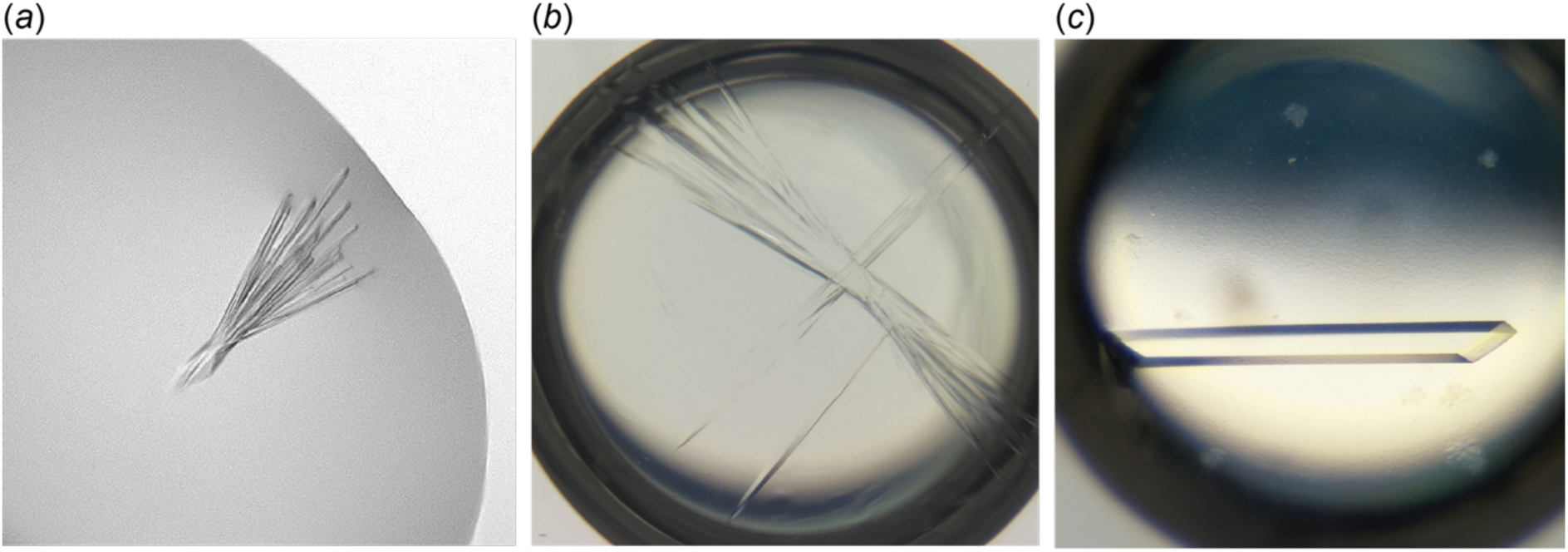
Progression of *Pm*UreE crystallisation from initial hit to seeded optimisation. (a) The initial sparse-matrix screen yielded a single crystallisation hit, resulting in thin needle crystals (Condition: 0.2 M ammonium citrate dibasic; 20% w/v polyethylene glycol (PEG) 3350). The image was recorded from a sitting-drop vapour-diffusion trial. (b) First-round optimisation around the initial condition generated longer needles (Condition: 0.14 M ammonium citrate dibasic, 18% w/v PEG 3350, with 100 μM NiCl_2_ added; a representative well is shown). The image was recorded from hanging-drop vapour-diffusion trials. (*c*) Microseeding using crushed microcrystals from the first-round optimisation markedly improved crystal quality, producing thicker, well-formed single crystals (Condition: 0.1 M sodium citrate, pH 5.0, 19% w/v PEG 6000). The image was recorded from hanging-drop vapour-diffusion trials.

**Figure S5.**
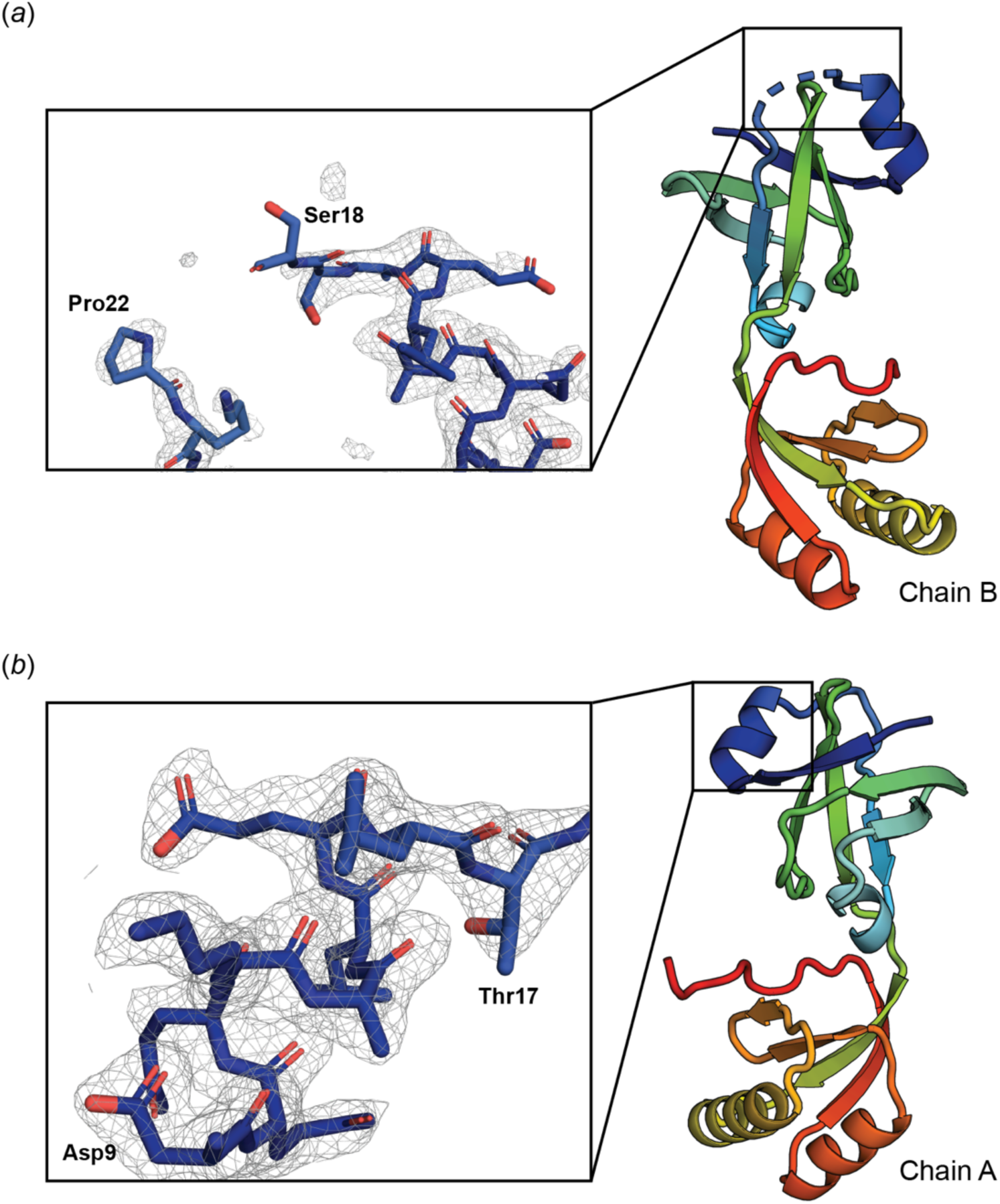
Key areas of electron density in the *Pm*UreE crystal structure. (*a*) The 2*m*F_o_-*D*F_c_ electron density map around the missing residues 19-21 in Chain B, contoured at 1.0 α. (*b*) The 2*m*F_o_-*D*F_c_ electron density map for the α1 helix in Chain A, contoured at 1.0 α, carve = 2.0. The maps were generated using phenix.mtz2map.

## References

Abrahams, G. & Newman, J. (2021). Acta Crystallogr D Struct Biol 77, 1437–1450.

Adams, P. D., Afonine, P. V., Bunkoczi, G., Chen, V. B., Davis, I. W., Echols, N., Headd, J. J., Hung, L. W., Kapral, G. J., Grosse-Kunstleve, R. W., McCoy, A. J., Moriarty, N. W., Oeffner, R., Read, R. J., Richardson, D. C., Richardson, J. S., Terwilliger, T. C. & Zwart, P. H. (2010). Acta Crystallogr D Biol Crystallogr 66, 213-221.

Afonine, P. V., Grosse-Kunstleve, R. W., Echols, N., Headd, J. J., Moriarty, N. W., Mustyakimov, M., Terwilliger, T. C., Urzhumtsev, A., Zwart, P. H. & Adams, P. D. (2012). Acta Crystallogr D Biol Crystallogr 68, 352–367.

Agirre, J., Atanasova, M., Bagdonas, H., Ballard, C. B., Basle, A., Beilsten-Edmands, J., Borges, R. J., Brown, D. G., Burgos-Marmol, J. J., Berrisford, J. M., Bond, P. S., Caballero, I., Catapano, L., Chojnowski, G., Cook, A. G., Cowtan, K. D., Croll, T. I., Debreczeni, J. E., Devenish, N. E., Dodson, E. J., Drevon, T. R., Emsley, P., Evans, G., Evans, P. R., Fando, M., Foadi, J., Fuentes-Montero, L., Garman, E. F., Gerstel, M., Gildea, R. J., Hatti, K., Hekkelman, M. L., Heuser, P., Hoh, S. W., Hough, M. A., Jenkins, H. T., Jimenez, E., Joosten, R. P., Keegan, R. M., Keep, N., Krissinel, E. B., Kolenko, P., Kovalevskiy, O., Lamzin, V. S., Lawson, D. M., Lebedev, A. A., Leslie, A. G. W., Lohkamp, B., Long, F., Maly, M., McCoy, A. J., McNicholas, S. J., Medina, A., Millan, C., Murray, J. W., Murshudov, G. N., Nicholls, R. A., Noble, M. E. M., Oeffner, R., Pannu, N. S., Parkhurst, J. M., Pearce, N., Pereira, J., Perrakis, A., Powell, H. R., Read, R. J., Rigden, D. J., Rochira, W., Sammito, M., Sanchez Rodriguez, F., Sheldrick, G. M., Shelley, K. L., Simkovic, F., Simpkin, A. J., Skubak, P., Sobolev, E., Steiner, R. A., Stevenson, K., Tews, I., Thomas, J. M. H., Thorn, A., Valls, J. T., Uski, V., Uson, I., Vagin, A., Velankar, S., Vollmar, M., Walden, H., Waterman, D., Wilson, K. S., Winn, M. D., Winter, G., Wojdyr, M. & Yamashita, K. (2023). Acta Crystallogr D Struct Biol 79, 449–461.

Armbruster, C. E., Mobley, H. L. T. & Pearson, M. M. (2018). EcoSal Plus 8.title>

Armbruster, C. E., Prenovost, K., Mobley, H. L. & Mody, L. (2017). J Am Geriatr Soc 65, 395-401.

Banaszak, K., Martin-Diaconescu, V., Bellucci, M., Zambelli, B., Rypniewski, W., Maroney, M. J. & Ciurli, S. (2012). Biochem J 441, 1017–1026.

Bellucci, M., Zambelli, B., Musiani, F., Turano, P. & Ciurli, S. (2009). Biochem J 422, 91–100.

Bergfors, T. (2007). Evolving Methods for Macromolecular Crystallography, edited by R. J. Read & J. L. Sussman: Springer.

Brauer, A. L., Learman, B. S. & Armbruster, C. E. (2020). Mol Microbiol 114, 185–199.

Brayman, T. G. & Hausinger, R. P. (1996). J Bacteriol 178, 5410-5416.

Chan, C. L., Pang, L. T. H., Choi, T., Chan, K. C., Tsang, K. L., Lee, K. M. & Wong, K. B. (2025). bioRxiv 2025.12.08.692910.

Colpas, G. J., Brayman, T. G., Ming, L. J. & Hausinger, R. P. (1999). Biochemistry 38, 4078–4088.

Colpas, G. J. & Hausinger, R. P. (2000). J Biol Chem 275, 10731–10737.

D’Arcy, A., Bergfors, T., Cowan-Jacob, S. W. & Marsh, M. (2014). Acta Crystallogr F Struct Biol Commun 70, 1117–1126.

Emsley, P., Lohkamp, B., Scott, W. G. & Cowtan, K. (2010). Acta Crystallogr D Biol Crystallogr 66, 486–501.

Evans, P. R. & Murshudov, G. N. (2013). Acta Crystallogr D Biol Crystallogr 69, 1204–1214.

Gouet, P., Courcelle, E., Stuart, D. I. & Metoz, F. (1999). Bioinformatics 15, 305–308.

Heimer, S. R. & Mobley, H. L. (2001). J Bacteriol 183, 1423-1433.

Higgins, D. G., Thompson, J. D. & Gibson, T. J. (1996). Methods Enzymol 266, 383–402.

Jones, B. D., Lockatell, C. V., Johnson, D. E., Warren, J. W. & Mobley, H. L. (1990). Infect Immun 58, 1120–1123.

Jumper, J., Evans, R., Pritzel, A., Green, T., Figurnov, M., Ronneberger, O., Tunyasuvunakool, K., Bates, R., Zidek, A., Potapenko, A., Bridgland, A., Meyer, C., Kohl, S. A. A., Ballard, A. J., Cowie, A., Romera-Paredes, B., Nikolov, S., Jain, R., Adler, J., Back, T., Petersen, S., Reiman, D., Clancy, E., Zielinski, M., Steinegger, M., Pacholska, M., Berghammer, T., Bodenstein, S., Silver, D., Vinyals, O., Senior, A. W., Kavukcuoglu, K., Kohli, P. & Hassabis, D. (2021). Nature 596, 583–589.

Kabsch, W. (2010). Acta Crystallogr D Biol Crystallogr 66, 125–132.

Kantardjieff, K. A. & Rupp, B. (2003). Protein Sci 12, 1865-1871.

Lee, M. H., Pankratz, H. S., Wang, S., Scott, R. A., Finnegan, M. G., Johnson, M. K., Ippolito, J. A., Christianson, D. W. & Hausinger, R. P. (1993). Protein Sci 2, 1042–1052.

Li, X., Zhao, H., Lockatell, C. V., Drachenberg, C. B., Johnson, D. E. & Mobley, H. L. (2002). Infect Immun 70, 389–394.

Liebschner, D., Afonine, P. V., Baker, M. L., Bunkoczi, G., Chen, V. B., Croll, T. I., Hintze, B., Hung, L. W., Jain, S., McCoy, A. J., Moriarty, N. W., Oeffner, R. D., Poon, B. K., Prisant, M. G., Read, R. J., Richardson, J. S., Richardson, D. C., Sammito, M. D., Sobolev, O. V., Stockwell, D. H., Terwilliger, T. C., Urzhumtsev, A. G., Videau, L. L., Williams, C. J. & Adams, P. D. (2019). Acta Crystallogr D Struct Biol 75, 861–877.

Matthews, B. W. (1968). J Mol Biol 33, 491–497.

McCoy, A. J., Grosse-Kunstleve, R. W., Adams, P. D., Winn, M. D., Storoni, L. C. & Read, R. J. (2007). J Appl Crystallogr 40, 658–674.

McPhillips, T. M., McPhillips, S. E., Chiu, H. J., Cohen, A. E., Deacon, A. M., Ellis, P. J., Garman, E., Gonzalez, A., Sauter, N. K., Phizackerley, R. P., Soltis, S. M. & Kuhn, P. (2002). J Synchrotron Radiat 9, 401–406.

Mobley, H. L. & Hausinger, R. P. (1989). Microbiol Rev 53, 85–108.

Mulrooney, S. B., Ward, S. K. & Hausinger, R. P. (2005). J Bacteriol 187, 3581–3585.

Nim, Y. S., Fong, I. Y. H., Deme, J., Tsang, K. L., Caesar, J., Johnson, S., Pang, L. T. H., Yuen, N. M. H., Ng, T. L. C., Choi, T., Wong, Y. Y. H., Lea, S. M. & Wong, K. B. (2023). Sci Adv 9, eadf7790.

Nim, Y. S. & Wong, K.-B. (2019). Inorganics 7.

Remaut, H., Safarov, N., Ciurli, S. & Van Beeumen, J. (2001). J Biol Chem 276, 49365–49370.

Sabbuba, N. A., Stickler, D. J., Mahenthiralingam, E., Painter, D. J., Parkin, J. & Feneley, R. C. (2004). J Urol 171, 1925–1928.

Schaffer, J. N., Norsworthy, A. N., Sun, T. T. & Pearson, M. M. (2016). Proc Natl Acad Sci U S A 113, 4494–4499.

Shi, R., Munger, C., Asinas, A., Benoit, S. L., Miller, E., Matte, A., Maier, R. J. & Cygler, M. (2010). Biochemistry 49, 7080–7088.

Song, H. K., Mulrooney, S. B., Huber, R. & Hausinger, R. P. (2001). J Biol Chem 276, 49359–49364.

Sriwanthana, B., Island, M. D., Maneval, D. & Mobley, H. L. (1994). J Bacteriol 176, 6836–6841.

Stickler, D. J. & Feneley, R. C. (2010). Spinal Cord 48, 784–790.

Stola, M., Musiani, F., Mangani, S., Turano, P., Safarov, N., Zambelli, B. & Ciurli, S. (2006). Biochemistry 45, 6495–6509.

Williams, C. J., Headd, J. J., Moriarty, N. W., Prisant, M. G., Videau, L. L., Deis, L. N., Verma, V., Keedy, D. A., Hintze, B. J., Chen, V. B., Jain, S., Lewis, S. M., Arendall, W. B., 3rd, Snoeyink, J., Adams, P. D., Lovell, S. C., Richardson, J. S. & Richardson, D. C. (2018). Protein Sci 27, 293-315.

Zambelli, B., Banaszak, K., Merloni, A., Kiliszek, A., Rypniewski, W. & Ciurli, S. (2013). J Biol Inorg Chem 18, 1005–1017.

Zeer-Wanklyn, C. J. & Zamble, D. B. (2017). Curr Opin Chem Biol 37, 80–88.

